# Oxidized phosphatidylcholines induce multiple functional defects in airway epithelial cells

**DOI:** 10.1101/823666

**Authors:** Christopher D Pascoe, Neilloy Roy, Emily Turner-Brannen, Alexander Schultz, Jignesh Vaghasiya, Amir Ravandi, Andrew J Halayko, Adrian R West

## Abstract

Oxidative stress is a hallmark of numerous airway diseases, contributing to extensive cell and tissue damage. Cell membranes and the airway mucosal lining are rich in phospholipids that are particularly susceptible to oxidative attack, producing bioactive molecules including oxidized phosphatidylcholines (OxPC). With the recent discovery of elevated OxPC in asthmatic patients after allergen challenge, we hypothesized that OxPC directly contribute to disease by inducing airway epithelial cell dysfunction.

We found that OxPC induced dose-dependent cell stress and loss of viability in BEAS-2B and Calu-3 cell lines and primary human epithelial cells. These responses corresponded with significant epithelial barrier dysfunction, which was further compounded when combining OxPC with an epithelial wound. OxPC inhibited DNA synthesis and migration required to re-establish barrier function, but cells recovered if OxPC were washed off soon after treatment. OxPC induced generation of reactive oxygen species, lipid peroxidation and mitochondrial dysfunction, raising the possibility that OxPC cause pathological lipid metabolism in a self-propagating cycle. The oxidative stress induced by OxPC could not be abrogated by putative OxPC receptor blockers, but partial recovery of barrier function, proliferation and lipid peroxidation could be achieved with the antioxidant n-acetyl cysteine.

In summary, we have identified OxPC as a group of bioactive molecules that significantly impair multiple facets of epithelial cell function, consistent with pathological features of asthma. Further characterisation of the mechanisms by which OxPC affect epithelial cells could yield new insights into how oxidative stress contributes to the pathogenesis of airway disease.

## INTRODUCTION

Asthma is a chronic disease of the airways which is characterized by inflammation, structural cell dysfunction, and excessive airway narrowing. Many of the features of asthma have been linked to reactive oxygen species (ROS) produced endogenously and upon exposure to environmental insults including air pollution, allergens, and smoke (49). ROS cause damage to tissues and cellular components and can act as a potent promoter of pro-inflammatory signals (45, 65, 66). Indeed, in airway epithelial cells, ROS enhances IL-8 production and reduces the activity of HDAC2 (30), which has been linked with steroid refractoriness in asthma (9, 37). Protein levels of glutathione peroxidase and superoxide dismuatase, two prominent antioxidant enzymes in the lung, are reduced in children with asthma (48). Further, the plasma levels of lipid peroxide markers of oxidative stress were increased in people with asthma, especially so in individuals with the glutathione-s-transferase val/val genotype (12). Finally, the abundance of ROS in the serum of patients with asthma is correlated with the degree of airway hyperresponsiveness (AHR) and acute exacerbations of asthma requiring hospitalization (8, 16).

ROS are capable of damaging lipid membranes and the lipid-rich mucosal lining of the airways resulting in the production of a group of potent pro-inflammatory molecules collectively known as oxidized phosphatidylcholines, or OxPC (4). OxPC are produced from the poly-unsaturated fatty acid tails of arachidonoyl and linoleoyl phospholipids to form OxPAPC and OxPLPC respectively (20). In disease, OxPC promote the induction of atherogenic pathways in endothelial cells (22, 35), and can modulate either enhancement or impairment of endothelial barrier function depending on the disease context (3, 33, 51). OxPC are also associated with inflammatory pain, mediated by transient receptor potential channels TRPA1 and TRPV1 (40). In the lung, OxPC have been identified in mice and humans following infection with influenza and *Mycobacterium* (10, 36, 55) and in the bronchoalveolar lavage fluid of patients with asthma (41). Some literature suggests that low doses of OxPC can act as inhibitors of toll-like receptor (TLR) signaling (13) however most research agrees that at pathophysiological concentrations, they act as mediators of inflammation and disease (21, 44). Recently, it has been shown in cultured human airway smooth muscle cells that OxPC promote inflammation characterized by IL-6, IL-8, GM-CSF secretion and the production of pro-inflammatory lipid mediators, and that OxPC induce airway narrowing in murine thin cut lung slices (41).

The airway epithelial cell plays an important role in protecting the underlying tissue from environmental irritants and coordinating inflammatory responses against microbes, and it is well established that dysfunction in the airway epithelium is an important factor asthma pathogenesis (28). With the recent identification of OxPC in the lungs of people with asthma and their well-established role in modulating endothelial cell function in blood vessels, we sought to establish their role in asthma pathogenesis through their ability to disrupt airway epithelial cell function. This manuscript is the first to show that OxPC promote airway epithelial cell dysfunction by disrupting their ability to form a protective barrier and impairing their self-repair capacity. Moreover, OxPC promote ongoing oxidative stress evidenced by the production of additional ROS and lipid peroxides, as well as mitochondrial dysfunction, consistent with the pathological features of asthmatic epithelium.

## METHODS

### Cell Culture

Calu-3 human airway epithelial cells were purchased from ATCC (HTB-55). Cells were propagated in DMEM/F-12 (Gibco 11330057) supplemented with 10% heat inactivated fetal bovine serum (FBS) (Gibco 12483020) and 1% penicillin/streptomycin (Gibco 15140122). For all experiments, Calu-3 were trypsinized and re-seeded at a density of 2×10^5^ cells/cm^2^ onto Transwell membranes [12 mm diameter, 0.4 μm pore size (Corning 3460)], or 18 mm round glass coverslips (VWR CA48382-041), 12-well plates (Sarstedt 83.3921), or Seahorse XF24 microplates (Agilent 100777-004) coated with 3 μg/mL rat tail collagen I (Corning 354236) in PBS. Cells were maintained in growth media for 4 days before being serum-deprived for 48 hours in DMEM/F12 containing 0.5% FBS and 1% insulin-transferrin-selenium (ITS) supplement (Gibco 41400045).

Normal human bronchial epithelial cells were purchased from Lonza (CC-2540S, single donor, lot # 493462) and grown in PneumaCult™-Ex Plus Medium (Stemcell Technologies 05040). Cells at passage 2-3 were trypsinized and re-seeded onto 12-well plates (nHBE) or Transwells (ALI-nHBE) at a density of 1×10^5^ cells/cm^2^ and maintained in PneumaCult™-Ex Plus for 3-4 days. ALI-nHBE cells were then transferred to fully supplemented PneumaCult™-ALI Medium, exposed to an air-liquid interface, and maintained for ≥3 weeks for mucociliary differentiation, confirmed by the presence of mucus, ciliary activity, and establishment of epithelial barrier function (>500 Ω.cm^2^).

BEAS-2B-GRE cells [gift from Dr. Rob Newton, previously described previously in (7)] were propagated in DMEM/F-12 supplemented with 10% FBS and 100 μg/mL G418, and seeded at a density of 3×10^3^ cells/cm^2^ in 12-well plates. Cells were grown until approximately 80% confluent then serum deprived for 24 hours in DMEM/F12 with 1% ITS.

### Generation and Administration of Oxidized Lipids

OxPC were generated as described previously (62). In brief, PAPC (1-palmitoyl-2-arachidonoyl-sn-glycero-3-phosphocholine, Avanti Polar Lipids 850459C) was exposed to room air for 3-4 days, during which PAPC visibly changed from a white to yellow paste. PAPC rapidly oxidizes in cell culture conditions, thus the non-oxidizable PSPC (1-palmitoyl-2-stearoyl-sn-glycero-3-phosphocholine, Avanti Polar Lipids 850456C) was used as a negative control . Treated OxPC/PSPC were resuspended in chloroform:methanol (2:1) at 1 mg/mL and stored at −20°C under nitrogen gas to prevent further oxidation.

OxPC and PSPC were prepared for dosing by aliquoting an appropriate volume of lipid solution and evaporating the chloroform:methanol under a stream of sterile nitrogen. The remaining lipid was immediately resuspended in warm media or buffer as appropriate and added to the cells (Calu-3 on Transwells: 1 mL basolateral chamber, 0.5 mL apical chamber; Calu-3, nHBE and BEAS-2B-GRE in 12-well plates: 1 mL; Calu-3 in Seahorse plates: 500 μL; ALI-nHBE on Transwells, 1 mL basolateral chamber, 75 μL apical chamber). For exposure times exceeding 3 days, cells were re-dosed with OxPC/PSPC in fresh media every 3-4 days. Apical OxPC and control treatments did not have any observable effect on the ciliary motion or mucus secretion of ALI-nHBE.

### Cell Stress and Viability

Cell stress was assessed by LDH release using the Roche Cytotoxicity Detection Kit (Roche 11644793001) in cells treated with OxPC (0-160 μg/mL) or PSPC (160 μg/mL) for 24 hours. Culture media was collected, gently centrifuged (150 × RCF for 4 minutes) and combined with the kit reagent for 10 min before absorbance readings were taken at 490 nm. LDH release was expressed as the percentage of LDH activity in the media relative to untreated control cells lysed with 2% triton X-100 (Sigma X-100).

Cell viability was determined by the uptake and retention of neutral red dye. After 24-hour exposure to OxPC or PSPC, a 0.33% stock solution of neutral red (Sigma N7005) in PBS was added 1:20 to the culture media and incubated for 2 hours. Cells were then gently rinsed with PBS and lysed in four times the original culture media volume with a solution of 1% acetic acid and 50% ethanol in H_2_O. Net absorbance was then calculated as A_540_ minus A_690_.

### Epithelial Barrier Function

The effect of OxPC on epithelial barrier function in Calu-3 and ALI-nHBE under static and scratched conditions was determined by measuring paracellular permeability and transepithelial electrical resistance (TEER). Paracellular permeability (*P*, cm/s) was calculated using the steady state approximation equation for Transwells (5):

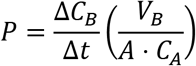

In brief, Transwells (surface area *A* = 1.13cm^2^) were treated with OxPC or PSPC for 24 hours, washed with PBS, and the apical chamber loaded with 0.5 mL of 300 μg/mL (*C*_*A*_) of sterile sodium fluorescein (NaFlu) (Sigma F6377) in Hank’s Balanced Salt Solution (HBSS) (Gibco 14065056). Assessment was initiated by transferring Transwells to fresh 12-well plates containing 1.5 mL (*V*_*B*_) per well HBSS, and monitoring NaFlu accumulation in the basolateral chamber at 37°C for 3 hours in a FLUOstar Optima fluorescence plate reader in orbital average mode (7 locations, 22 mm diameter, Ex: 460 nm Em: 520 nm). Fluorescence time curves for each Transwell were converted to basolateral concentration (*C*_*B*_, μg/mL) time curves using a fluorescence-concentration standard curve. The slope of the linear segment of these concentration-time curves (*ΔC*_*B*_/*Δt*, μg/mL/s) was then determined using the linear regression function of GraphPad Prism 6.07.

TEER was measured with a freshly calibrated MilliCell ERS-02 Voltohmmeter (Millipore). Baseline TEER was measured in PBS prior to cell seeding and subtracted from all experimental measurements, and net TEER was normalized to an equivalent untreated and non-scratched control to account for temporal variability. To assess barrier recovery after a wound, cells were scratched with a sterile 10 μL pipet tip and firm pressure, washed once with PBS, and post-wound TEER readings taken. Cells were then treated with media containing OxPC or PSPC as appropriate, and TEER monitored for up to 7 days.

### DNA synthesis as a metric of proliferation

DNA synthesis was assessed by thymidine-analog incorporation and histological detection using the Click-iT EdU Imaging Kit (Thermo Fisher C10337). Confluent serum-deprived Calu-3 cells on coverslips were scratched, washed, and incubated with OxPC (80 μg/mL) for up to 48 hours before being pulsed with 10 μM 5-ethynyl-2’-deoxyuridine (EdU) for 2 hours. The cells were then fixed with 4% paraformaldehyde (PFA) in PBS for 15 minutes, washed twice with 3% BSA in PBS and permeabilized with 0.5% triton X-100 in PBS for 20 minutes. EdU fluorescent labelling was performed per the manufacturers instruction’s before nuclei were counterstained with 0.2 μg/mL Hoechst 33342 (Thermo Fisher H1399) in PBS for 10 minutes. Coverslips were mounted with SlowFade Diamond (Thermo Fisher S36967) before being sealed with nail varnish and imaged on an Olympus IX-51 fluorescent microscope. Images of at least 3 fields from each coverslip (minimum 1000 cells per ‘n’) were captured at the same time with consistent exposure settings, before EdU positive and total nuclei were manually counted with ImageJ.

### mRNA abundance

The mRNA abundance for epithelial phenotype markers was assessed by RT-qPCR. Calu-3 were treated with OxPC or PSPC for 18 hours before total RNA was isolated by the Ambion Purelink RNA Mini Kit (Thermo Fisher 12183025) before RNA concentration and purity were assessed by A_260_:A_280_ spectrophotometry. Reverse transcription was performed with 1 μg total RNA using iScript Reverse Transcription Supermix for RT-qPCR (Bio-Rad 170-8841). cDNA equivalent to 20 ng of total RNA was then amplified in duplicate using SsoAdvanced Universal SYBR Green Supermix (Bio-Rad 172-5274) and 300 nM of the appropriate primers (Table 1) in a Bio-Rad CFX96 thermal cycler. Cycling conditions involved 30 s incubation at 95°C for polymerase activation, followed by 40 cycles of denaturing at 95°C for 10 s and annealing/extension at 58°C for 30 s. Crossing thresholds (Ct) were calculated and housekeeping gene stability confirmed (M value <0.25) using the Bio-Rad CFX Manager 3.1 software, and mean reaction efficiency (E) was determined using LinRegPCR 2017.0 (46). Relative mRNA abundance was calculated using efficiency-corrected ΔCt (E^ΔCt^) divided by a normalizing factor (NF) derived from the geometric mean of E^ΔCt^ for no less than two stable housekeeping genes (26, 56). The final data was scaled such that untreated control condition presented a mean result of 1 AU.

**TABLE 1.**
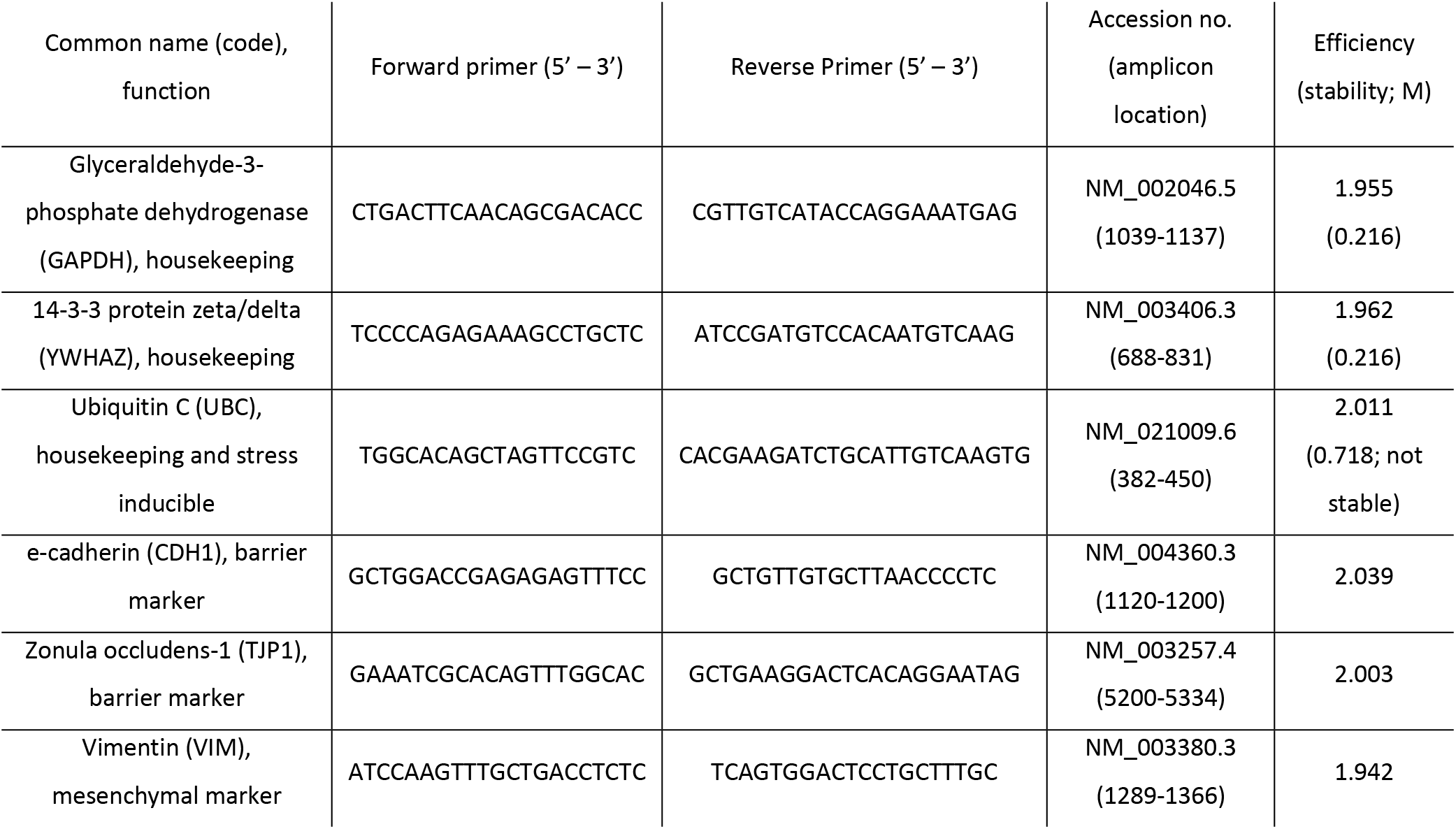
Primers used for mRNA abundance analysis by RT-qPCR. Primers were selected using Primer-BLAST [http://www.ncbi.nlm.nih.gov/tools/primer-blast/ (61)] to span exon-exon junctions to prevent amplification of genomic DNA, with primer specificity continually assessed by melting curve analysis.

### Oxidative Stress

The capacity of OxPC and house dust mite extract (HDM, Greer XPB82D3A2.5) to induce the accumulation of general cytosolic ROS, cytosolic superoxide and mitochondrial superoxide in Calu-3 were assessed by 2′,7′-dichlorofluorescin diacetate (DCF), dihydroethidium (DHE) and MitoSOX assays respectively. For DCF assays, serum-deprived Calu-3 were loaded with 10μM DCF (Sigma D6883) in low-serum media for 30 minutes, then washed with warm Hanks Balanced Salt Solution (HBSS). Cells were then incubated with OxPC, PSPC, HDM (100 μg/mL) or positive control cumene hydroperoxide (cumene-OOH, 200 μM) in HBSS, and fluorescence quantified in a FLUOstar Optima (Ex 492, Em 520) for 2 hours.

For DHE and MitoSOX assays, cells were pre-incubated with OxPC, PSPC, or positive controls doxorubicin (50 μM; New England Biolabs 5927) and antimycin A (10 μM; Sigma-Aldrich A8674) for 2 hours in low-serum media. Cells were then stained with 5μM DHE (Thermo Fisher D1168) for 30 minutes or 5μM MitoSOX (Thermo Fisher M36008) for 15 minutes and counterstained with H33342 for 10 minutes. Fluorescence intensity was quantified in a FLUOstar Optima (DHE and MitoSOX: Ex 485, Em 590), and cells were imaged by fluorescence microscopy. Exposure was set using the appropriate positive control condition, and all images were taken at the same time at consistent settings.

### Lipid Peroxidation Assays

The generation of lipid peroxides in response to OxPC- and HDM-induced-ROS was quantitatively assessed in Calu-3 with the Image-iT Lipid Peroxidation Kit (Thermo Fisher C10445). Calu-3 cells were loaded with 10 μM BODIPY-C11 in HBSS for 30 minutes, washed three times with PBS, and treated with OxPC, PSPC, HDM (100 μg/mL) or cumene-OOH in HBSS for 2 hr. The progress of lipid peroxidation was monitored as the ratio of oxidized to reduced BODIPY-C11 at 37°C for 2 hours in a Cytation 5MPV (fluorescence intensity 5×5 area scanning, Oxidized Ex: 478/20 nm Em 520/20 nm, Reduced Ex: 566/20 nm Em 606/20 nm).

To confirm lipid peroxidation and subsequent protein oxidization, the Click-iT Lipid Peroxidation Imaging Kit (Thermo Fisher C10446) was used. Calu-3 cells were simultaneously loaded with 50 μM linoleamide alkyne (LAA) and treated with OxPC in HBSS for 2 hours. Cells were then washed with PBS, fixed in 4% PFA for 15 minutes then permeabilized in 4% PFA with 0.5% triton X-100. After 3 washes in PBS cells were blocked with 1% BSA for 30 minutes and LAA fluorescent labelling was performed per the manufacturer’s instructions. Cells were then counterstained, mounted and imaged as described above.

### Mitochondrial Function

Mitochondrial membrane potential was assessed in live cells using the fluorescent dye tetramethylrhodamine (TMRM; Biotium 70017). Briefly, Calu-3 were treated with OxPC or PSPC for 2-hours and then stained with 0.1 μM TMRM in HBSS for 30-minutes. The oxidative phosphorylation uncoupler CCCP (2 μM, Sigma C2759) was used as a negative control to establish baseline/minimal mitochondrial membrane potential. Excess TMRM dye was removed with HBSS and the degree of fluorescence intensity relative to the CCCP condition was measured in a FLUOstar Optima (Ex 548, Em 573 nm).

Oxygen consumption rate (OCR) was measured during Seahorse XF Cell Mito Stress Tests as per the user guide (Agilent 103015-100) with modifications. In brief, Calu-3 in Seahorse plates were switched to Seahorse XF base media (Agilent 103334-100) supplemented with 10 mM glucose (Fisher BP350), 1 mM sodium pyruvate (Gibco 11360070) and 2 mM Glutamax (Gibco 35050061) and treated with OxPC such that the first baseline OCR reading was timed 2 hr after exposure. Seahorse Sensor Cartridges (Agilent 100867-100) were hydrated for 6-18 hours at 37 °C in 1 mL of the supplied calibrant. Stock solutions of stress test reagents were prepared by dissolving in DMSO, diluting to 10× desired final concentrations with glucose/pyruvate/Glutamax supplemented media, before adding to the appropriate sensor ports. The sensor cartridge was calibrated in a Seahorse XF24 Analyzer, cell plates installed, and OCR measured for 30 min at baseline, and in response to 2μM oligomycin (New England Biolabs 9996L), 0.5 μM FCCP (Sigma C2920), and 0.5 μM rotenone/antimycin-A (Sigma R8875 and A8674 respectively). Triplicate technical replicates on each plate were averaged to produce final OCR datapoints, and parameters of mitochondrial function calculated according to the user guide.

### Prevention of OxPC induced dysfunction

ROS formation was used as an initial parameter to determine whether dysfunction induced by OxPC could be prevented. Calu-3 were pre-incubated for 2 hours with vehicle control (0.1% DMSO), inhibitors of ROS [N-acetyl-L-cysteine (NAC), Sigma-Aldrich A7250], or blockers of receptors associated with OxPC activity including platelet activating factor receptor (WEB-2086, Tocris 2339), prostaglandin EP2 receptor (PF-04418948, Tocris 4814), CD36 [sulfosuccinimidyl oleate (SSO), Cayman Chemical 11211] and toll like receptors 2 and 4 [sparstolonin B (SsnB), Sigma-Aldrich SML1767]. Cells were then loaded with DCF and co-incubated with inhibitors, OxPC (160 μg/mL) and vehicle control (0.1% DMSO) for a further 2 hours before measuring fluorescence. Subsequently, Calu-3 were treated concurrently with OxPC (80 or 160 μg/mL) and NAC (0.1 to 10 mM) before being subjected to TEER, scratch/proliferation and lipid peroxidation assays.

### Data Analysis and Statistics

All numerical data are presented as mean ± standard error. Each ‘n’ value represents a fully independent experiment created from a separate flask of stock cells. Statistical tests were performed with the GraphPad Prism 6.07 software package, with p<0.05 considered statistically significant. Matched samples 1-way ANOVA was used for all OxPC experiments where data was collected at a single time-point, with Dunnett post-tests calculated against control or OxPC 160 μg/mL conditions as appropriate. In TEER recovery experiments, all pairwise Bonferroni post-tests were calculated. For time-course experiments, independent samples 2-way ANOVA (factors: OxPC, time) was performed, with Dunnett post-tests calculated against the untreated control condition. Paired t-test was used for all HDM experiments.

## RESULTS

### OxPC induce cell stress and loss of viability in airway epithelial cells

OxPC treatment resulted in dose-dependent LDH release, indicating cell stress in each cell type tested (Fig 1A). However, the magnitude of the effect was variable across epithelial cell lines with strong responses at low doses in BEAS-2B (p<0.0001), moderate responses in Calu-3 (p<0.0001), and weak responses in nHBE (p<0.0001) and ALI-nHBE (p=0.0257). Very high levels of LDH release were associated with loss of viability in BEAS-2B (Fig 1B; p<0.0001) and Calu-3 (p<0.0001) cells. No loss of viability was detected in nHBE (p=0.2204). Statistical significance was achieved in ALI-nHBE (p=0.0438), however the magnitude of any effect was small relative to the other cell types, with no obvious dose-dependence and no significant differences relative to control in post-tests. The non-oxidizable negative control PSPC caused no quantitative or qualitative changes in this experiment, or in any subsequent experiment.

**Figure 1.**
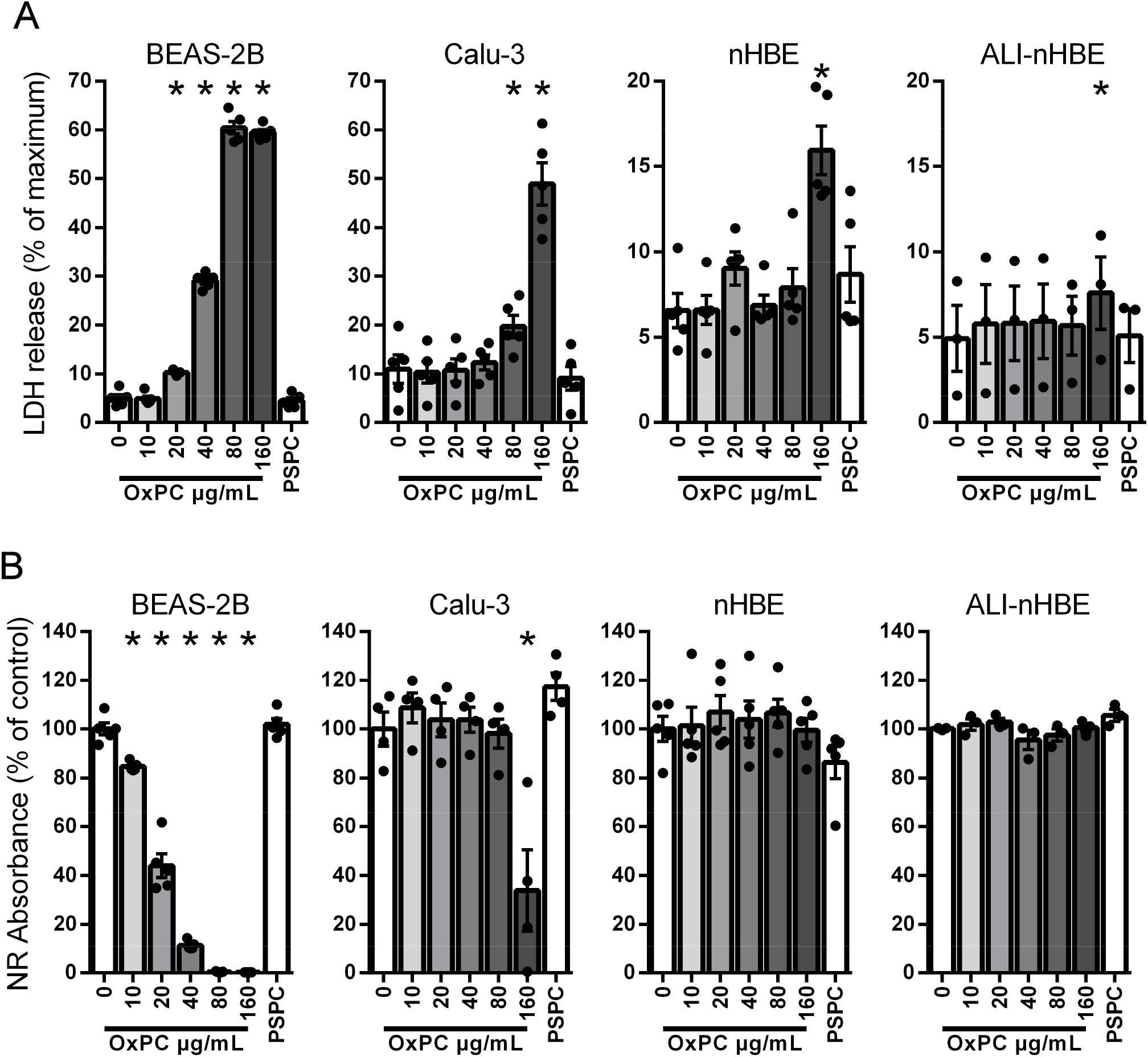
OxPC induces cell stress (A; increased LDH release) and loss of cell viability (B; reduced NR absorbance) in a dose-dependent and cell-specific manner. BEAS-2B were particularly sensitive experiencing almost complete cell death at 40 μg/mL OxPC (stress p<0.0001, viability p<0.0001, n=5). Calu-3 were moderately sensitive (stress p<0.0001, viability p<0.0001, n≥4) with cytotoxicity only apparent at 160 μg/mL OxPC, whereas nHBE (stress p<0.0001, viability p=0.2204, n=5) and ALI-nHBE (stress p=0.0257, viability p=0.0438, n=3) were relatively robust. * represents a significant difference from the 0 μg/mL OxPC control condition.

### OxPC reversibly impair epithelial barrier function and barrier recovery after a wound

In Calu-3 cells, OxPC exposure at 80 and 160 μg/mL resulted in significant barrier impairment, as measured by both paracellular permeability (Fig 2A; p<0.0001) and TEER assays (Fig 2B; OxPC p<0.0001). Large voids in the monolayer representing cell loss were evident after 24 hr in 160 μg/mL OxPC but no cell loss was apparent at 80 μg/mL even after 72 hours, consistent with cell stress and viability assays. The 40 μg/mL dose demonstrated enhanced barrier function by TEER at 24 hours, but this effect was not significant from 48 hours onwards. For ALI-nHBE, OxPC treatment did not cause any difference in permeability (Fig 2C; p=0.5510), but resulted in significant impairment of TEER (Fig 2D; OxPC p=0.0007), with strong dose-dependent dysfunction at seven days.

**Figure 2.**
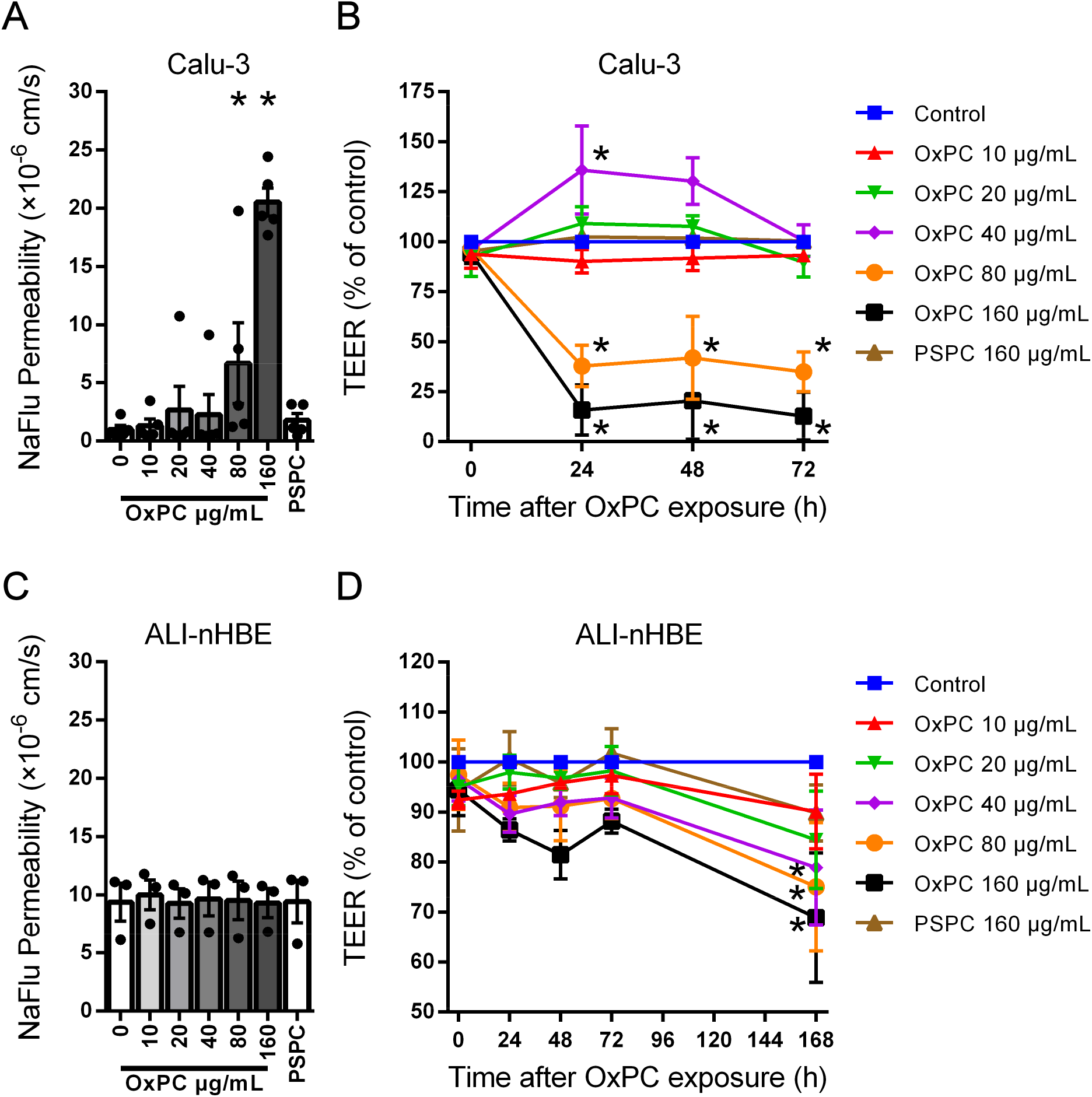
Under static conditions, barrier function was impaired by OxPC in a dose-dependent and cell-specific manner. Calu-3 exhibited dysfunction in both paracellular permeability (A; p<0.0001, n=5) and TEER (B; interaction p<0.0001, time p=0.0030, OxPC p<0.0001, n=4) at 80-160 μg/mL. The effect of OxPC on ALI-nHBE is not significant for permeability (C; p=0.5510, n=3) but is significant for TEER (D; interaction p=0.9909, time p=0.0007, OxPC p=0.0232, n=3), with strong dose dependent dysfunction apparent at 7 days. * represents difference from the 0 μg/mL OxPC control condition.

Combining OxPC with a physical scratch revealed that the simultaneous ‘two hits’ could cause additional barrier dysfunction in Calu-3 cells (Fig 3A; OxPC p<0.0001). TEER was used to assess wound healing, as it collectively represents the migration, proliferation and junction formation essential to re-establish barrier function. Consistent with unscratched cultures, the 80 and 160 μg/mL groups displayed a failure of wound healing after 24 hours. Surprisingly, combining OxPC at 40 μg/mL with a scratch resulted in a significant delay in wound healing at 24-48 hours, in stark contrast to the barrier enhancement seen with OxPC treatment alone. To determine whether the effects of OxPC on barrier function after a wound are reversible, scratched Calu-3 were exposed to OxPC for 24 or 72 hours, before the OxPC were washed off, and TEER re-measured after 24 hours. Only OxPC 80 μg/mL treated cells could partially recover barrier function (Fig 3B; p=0.0079) and the recovery was more complete with shorter pre-exposure times.

**Figure 3.**
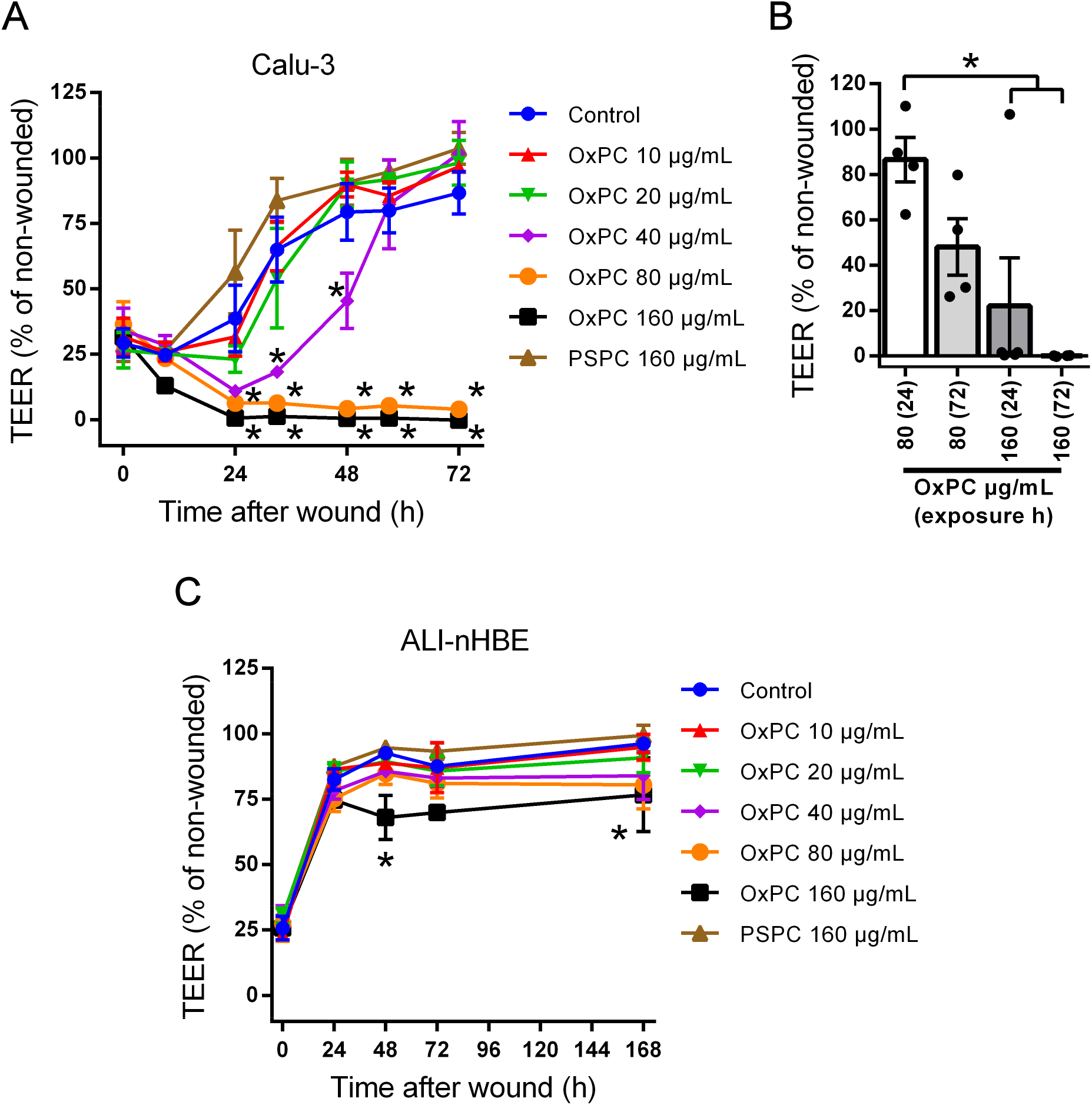
OxPC treatment potentiates barrier dysfunction after a wound. In Calu-3 cells, OxPC at 40 μg/mL significantly delayed recovery of barrier function (contrast with the lack of defect at 40 μg/mL in Fig 2B) with ongoing dysfunction evident at 80-160 μg/mL (A; interaction p<0.0001, time p<0.0001, OxPC p<0.0001, n≥4). However, barrier function could partially recover in the 80 μg/mL condition if OxPC are rapidly washed off (B; p=0.0079n≥4). Scratched ALI-nHBE cultures display defective wound healing in the OxPC 160 μg/mL group (C; interaction p=0.8773, time p<0.0001, OxPC p<0.0001) that is not evident until later timepoints in static cultures (contrast with the lack of significant defect for 160 μg/mL at 48 hrs in Fig 2D). * represents significant difference from the 0 μg/mL OxPC control condition (wound healing) or between conditions (washout recovery).

In ALI-nHBE, OxPC at 160 μg/mL caused a modest but highly significant impairment of wound healing at 48 hours, with dose-dependent dysfunction persisting until 7 days (Fig 3C; OxPC p<0.0001).

### OxPC reversibly inhibit DNA synthesis

Airway epithelial wound healing involves an initial migratory phase followed by increased proliferation at the wounded edge (63), thus we histologically assessed DNA synthesis to better understand the dynamics of barrier recovery in the presence of OxPC (Fig 4). Scratched cultures exhibit increased EdU labelling indicative of DNA synthesis at the leading edge. In contrast, a submaximal 80 μg/mL OxPC dose appropriate for multi-day experiments resulted in a dramatic decrease in DNA synthesis at 24 hours (p<0.0001) DNA synthesis returns to control levels if OxPC are left on the cells for 48 hours but experiences a dramatic rebound if OxPC are washed off the cells, albeit without immediate wound closure. These results are consistent with the barrier function results following both scratch and wash protocols.

**Figure 4.**
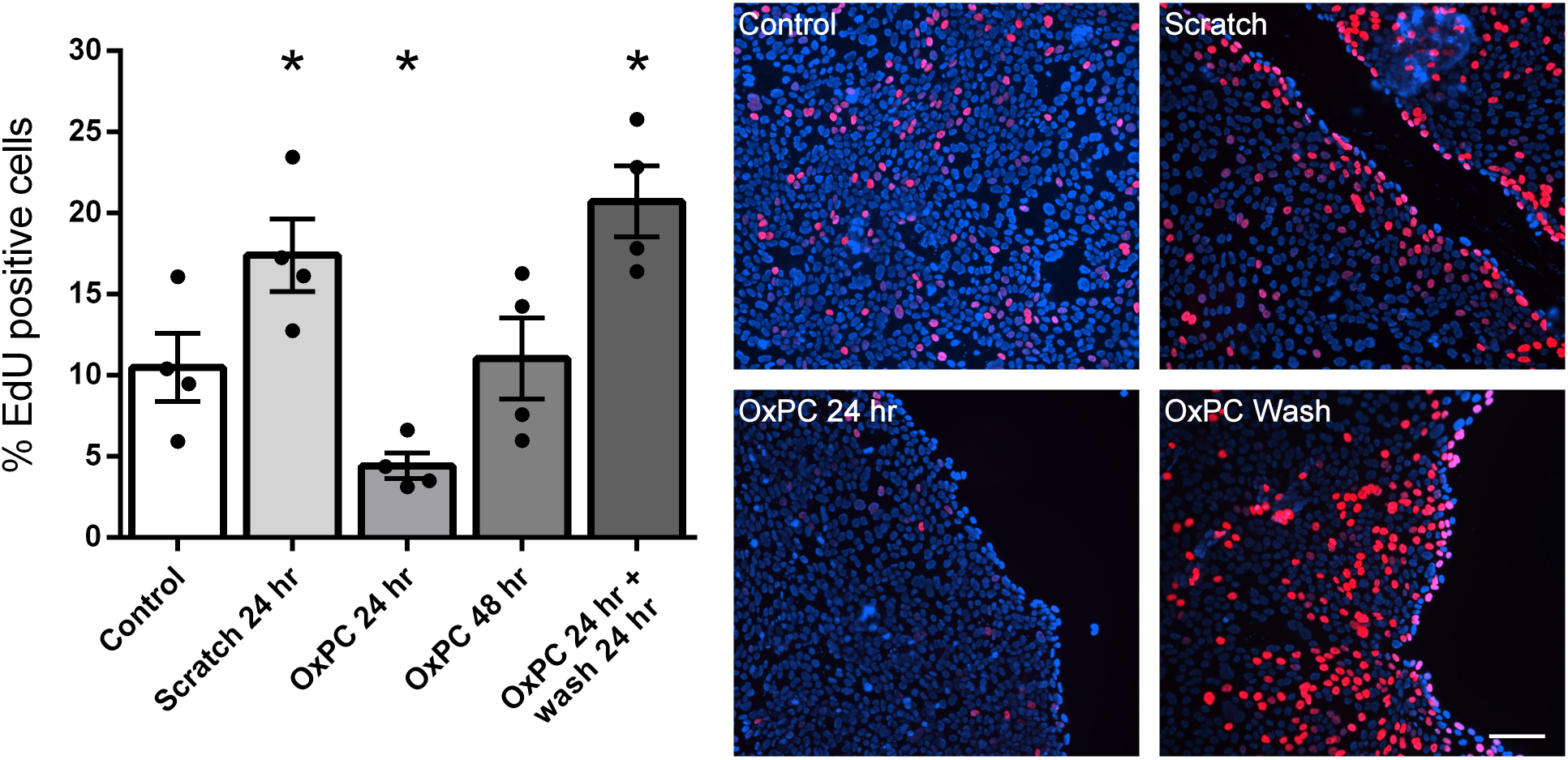
DNA synthesis in Calu-3 cells is upregulated on the leading edge after a scratch, but this is significantly inhibited by 80 μg/mL OxPC (p<0.0001, n=4). DNA synthesis recovers if OxPC are washed off, but without immediate wound closure. * represents significant difference from the 0 μg/mL OxPC control condition. Scale bar is 100 μm.

### OxPC do not affect epithelial phenotype markers

To determine whether barrier dysfunction was modulated by changes in the epithelial phenotype, the relative mRNA abundance of three key phenotypic markers was measured after OxPC exposure (Fig 5). Neither e-cadherin (CDH1, p=0.4295) nor vimentin (VIM, p=0.4362) mRNA levels were altered by OxPC, suggesting maintenance of epithelial phenotype and no acquisition of mesenchymal features. Zonula occludens 1 (TJP1) mRNA levels were significantly different across the groups in the ANOVA statistic (p=0.0069), however there was no linear dose-dependence and post-tests revealed no individual group differences. Intriguingly, ubiquitin C (UBC) was dose-dependently regulated by OxPC (p<0.0001). This gene was intended for usage as a housekeeping gene as described previously (58) but is also known to be stress-inducible (19), supporting that OxPC induce cellular stress in epithelial cells.

**Figure 5.**
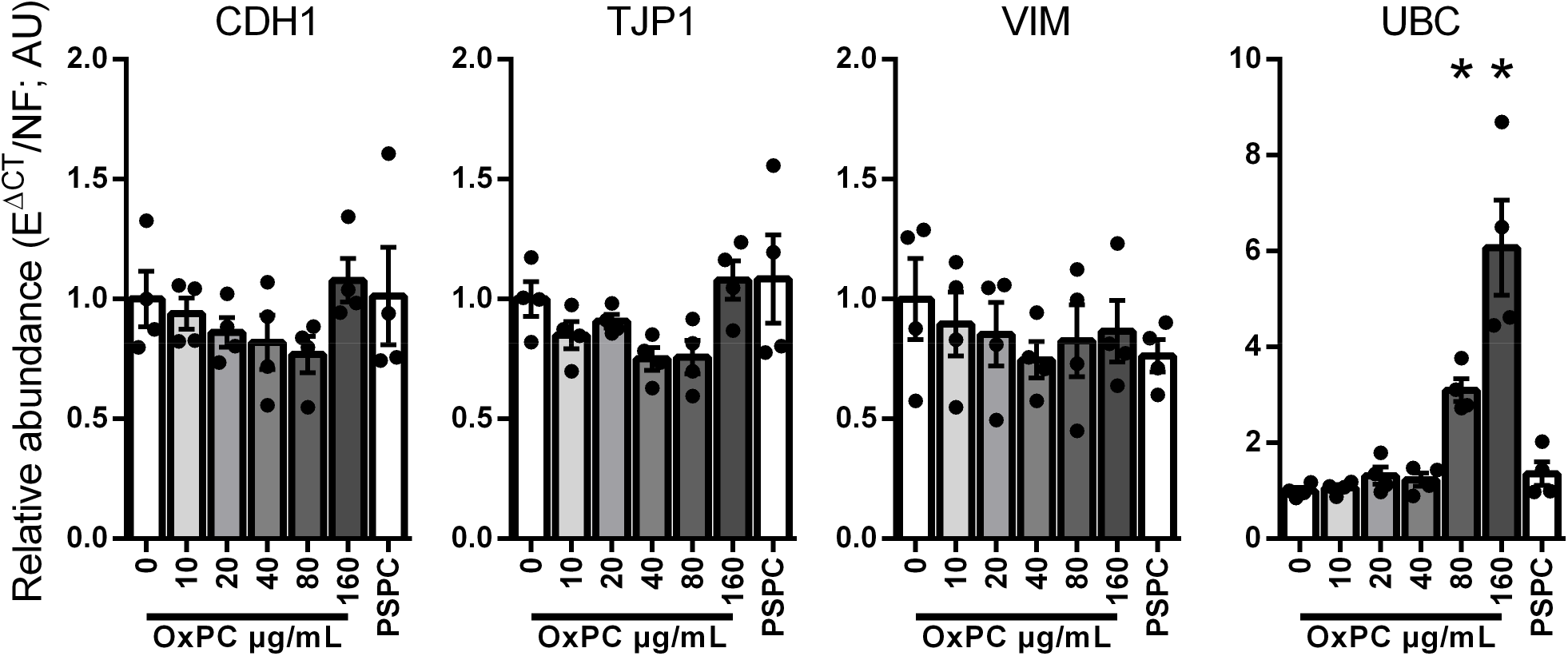
Markers for epithelial and barrier phenotype are not appreciably regulated by OxPC in Calu-3 cells. The mRNA abundance of CDH1 (p=0.4295, n=4) and VIM (p=0.4362, n=4) were not significantly altered after OxPC treatment. While statistical significance was detected for TJP1 mRNA levels (p=0.0069, n=4), the small magnitude of the changes and lack of dose-dependency makes it unlikely that modulation of this gene is responsible for the observed barrier dysfunction. In contrast, the stress-inducible gene UBC demonstrated strong dose-dependent regulation by OxPC (p<0.0001, n=4).

### OxPC induce general ROS accumulation

The established relationship between OxPC and ROS, combined with our collective cell stress data, led us to examine the effect of OxPC on ROS accumulation. OxPC rapidly induced general ROS accumulation in a time- and dose-dependent manner (Fig 6A-B; p<0.0001). To put these results in context of known disease-linked modulators, we performed separate experiments exposing cells to a maximal dose of house dust mite (HDM). General ROS accumulation was significantly elevated in response to HDM (p=0.0314), although the magnitude was markedly lower than a maximal OxPC dose.

**Figure 6.**
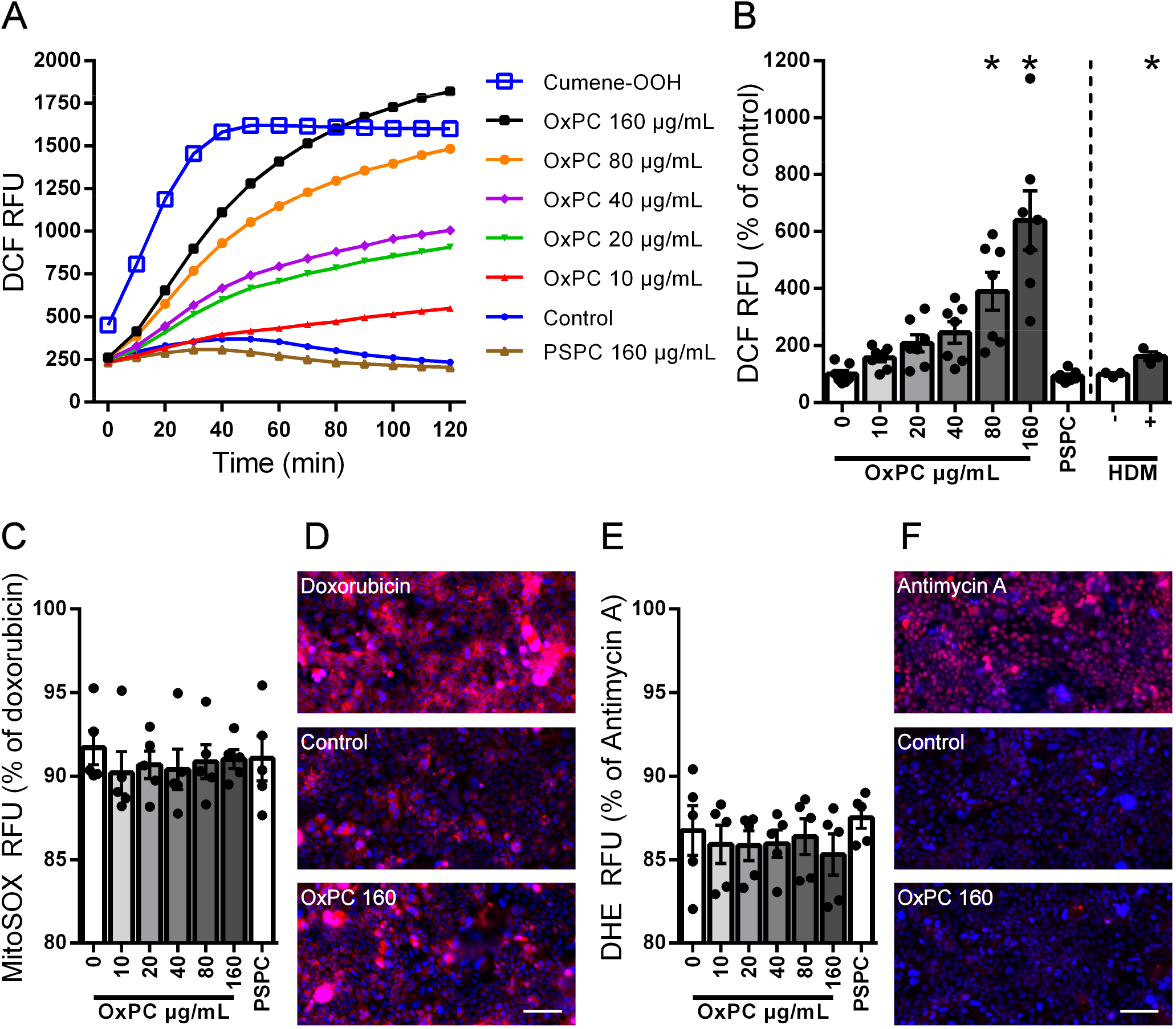
OxPC induce time- (A) and dose-dependent (B) general ROS accumulation in Calu-3 cells (p<0.0001, n=7). HDM is also capable of inducing ROS accumulation (p=0.0314, n=3), but with a much smaller magnitude. Mitochondrial (C-D; p=0.3730, n=5) and cytosolic (E-F; p=0.0227, n=5) superoxide are only marginally influenced by OxPC, with no obvious dose-dependence and significantly lower signal than their respective positive controls. * represents significant difference from the 0 μg/mL OxPC control condition. Scale bar is 100 μm.

To try and ascertain the specific cellular source of ROS, we assessed the accumulation of mitochondrial and cytosolic superoxide. Mitochondrial superoxide was not significantly altered by OxPC treatment (Fig 6C; p=0.3730). Background fluorescence levels were high for this assay when quantified with a plate reader, but no quantitative or qualitative difference was apparent between control and OxPC conditions when compared with the brightly stained positive control (immunofluorescence microscopy; Fig 6D). Cytosolic superoxide reached statistical significance (Fig 6E; p=0.0227) however there was no obvious dose-dependence, post-tests were not significant, and the overall magnitude of any OxPC-related response was considerably lower than the corresponding positive control. Moreover, fluorescent imaging shows no qualitative difference between the control and maximal dose of OxPC relative to the much brighter positive control (Fig 6F). These combined results indicate that superoxide accumulation is unlikely to be biologically significant during OxPC exposure.

### OxPC induce additional lipid peroxidation

To determine whether ROS formation could induce ongoing cellular dysfunction, we assessed lipid peroxide generation. Similar to general ROS accumulation, lipid peroxides began accumulating rapidly after OxPC addition, with higher doses surpassing the level of lipid peroxides produced by the positive control cumene-OOH within 60 minutes (Fig 7A). At the 2-hour endpoint, all OxPC doses were significantly higher than control (Fig 8B; p<0.0001), with a strong dose-dependency. In separate experiments, HDM was assessed for its ability to induce lipid peroxidation. The effect of HDM approached statistical significance (p=0.0522), but the magnitude of the change was smaller than even the lowest OxPC dose.

**Figure 7.**
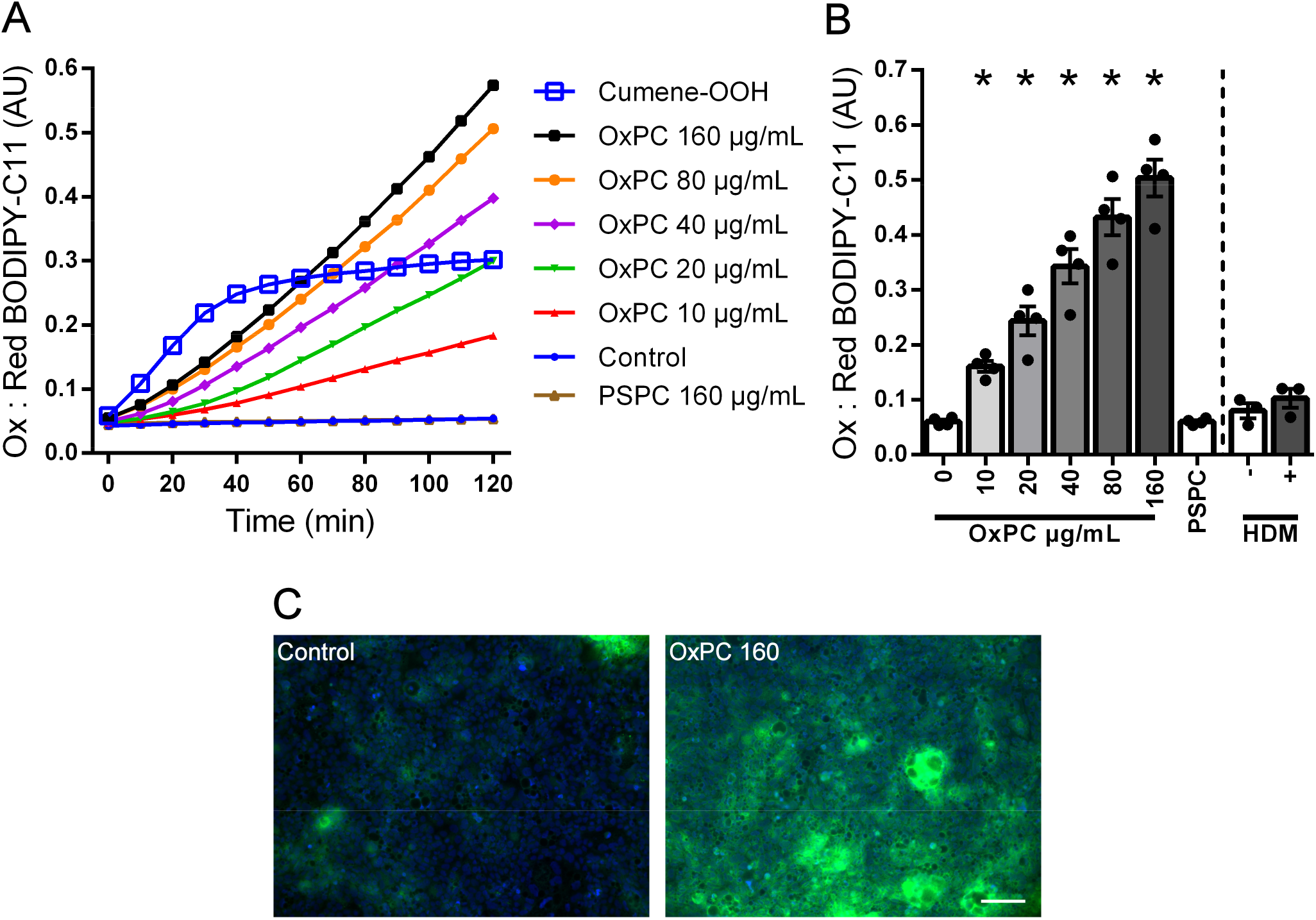
OxPC induce time- (A) and dose-dependent (B) generation of lipid peroxides in Calu-3 cells (p<0.0001, n=4). The effect of HDM was modest in comparison and not statistically significant (p=0.0522, n=3). In contrast to general ROS, 10 mM NAC could only partially reverse lipid peroxidation (C; p=0.0001, n=7). Assessing lipid peroxidation by an alternative histological technique (D) confirms significant accumulation that did not appear to be localized to any specific cell membrane or organelle. * represents significant difference from the 0 μg/mL OxPC control condition (B) and from the OxPC 160 μg/mL condition (C). Scale bar is 100 μm.

**Figure 8.**
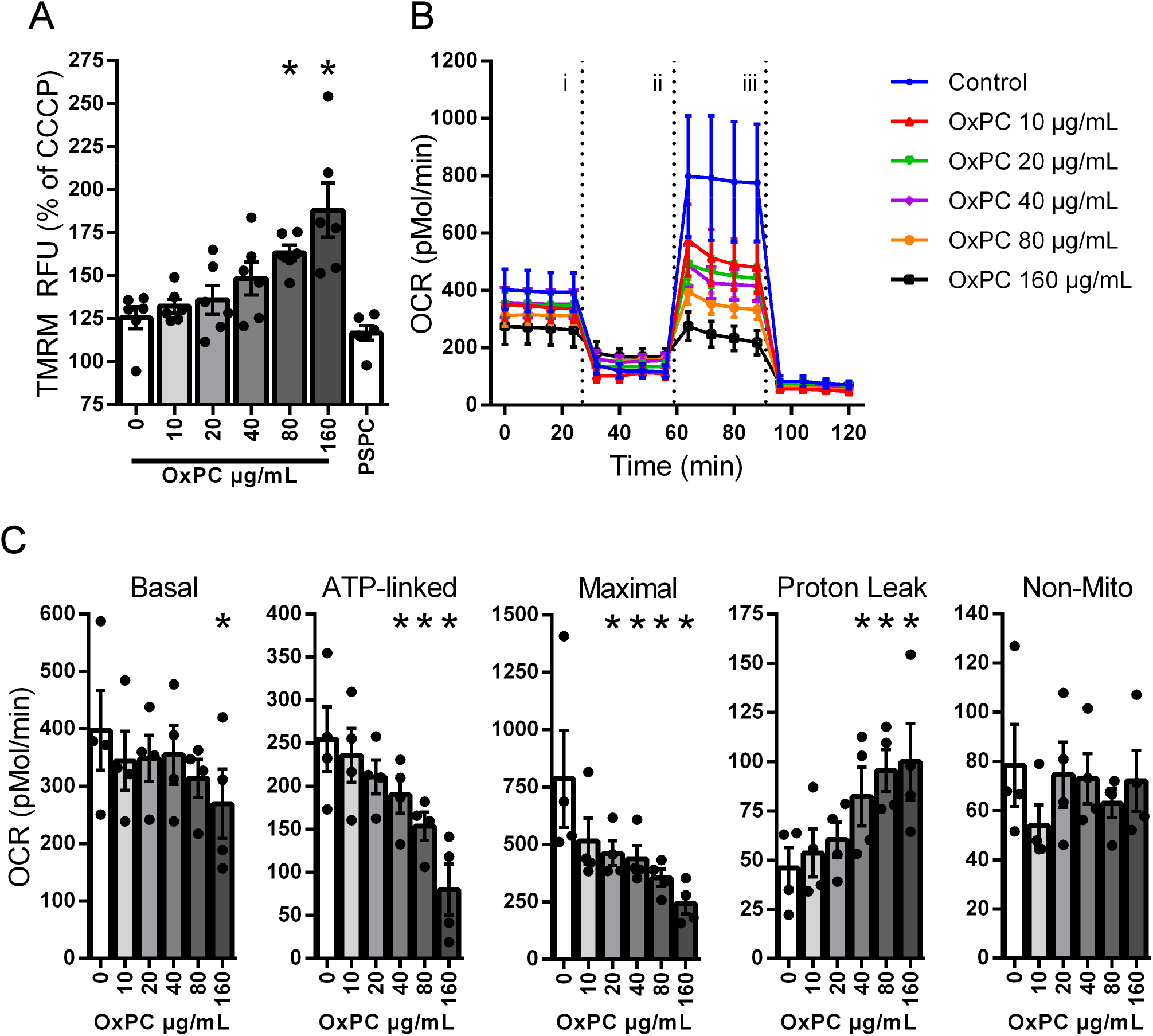
OxPC induce dose-dependent increase in TMRM fluorescence, indicative of mitochondrial membrane hyperpolarisation (A; p<0.0001, n=6). Reductions in OCR (interaction p=0.0001, time p<0.0001, OxPC p<0.0001, n=4) are apparent during mitochondrial stress tests (B; i=oligomycin, ii=FCCP, iii=rotenone/antimycin-A), and calculation of distinct metabolic parameters (C) reveals dose-dependent reductions in basal (p=0.0165), ATP-linked (p<0.0001) and maximal respiration (p=0.0013), dose-dependent increases in proton leak (p=0.0002), but no change in non-mitochondrial respiration (p=0.0952). These combined responses are indicative of mitochondrial dysfunction in response to OxPC challenge. * represents significant difference from the 0 μg/mL OxPC control condition.

To validate these results, lipid peroxidation was assessed by an alternative histological technique that assesses lipid peroxide transfer to proteins (Click-iT Lipid Peroxidation Imaging Kit). OxPC induced a qualitative increase in staining indicating lipid peroxidation and subsequent protein peroxidation, that was diffuse across the cell membrane and cytoplasm. There was no obvious segregation or punctate patterning that would indicate isolation to any specific organelle (Fig 8D).

### Mitochondrial function is disrupted by OxPC treatment

Oxidized phospholipids accumulate in the mitochondria (62), and oxidized lipoproteins can cause mitochondrial membrane hyperpolarisation (23). Moreover, mitochondria are a well-known source of multiple ROS species, and mitochondrial metabolism is required for proliferation. Thus, we determined the effect of OxPC on Calu-3 mitochondrial membrane potential, as well as oxygen consumption during a mitochondrial stress test. Consistent with earlier work, OxPC induced a dose-dependent increase in TMRM staining, indicative of mitochondrial hyperpolarisation (Fig 8A; p<0.0001). Moreover, OxPC caused profound alterations in oxygen consumption rate (OCR) during the stress test (Fig 8B; OxPC p<0.0001), with specific dose-dependent reductions in each of basal respiration (Fig 8C; p=0.0165), ATP-linked respiration (p<0.0001) and maximal respiration (p=0.0013), dose-dependent increases in proton leak (p=0.0002), but no change in non-mitochondrial respiration (p=0.0952).

### OxPC receptor blockade does not mitigate OxPC-induced ROS accumulation

In other cell types, the negative effects of OxPC are associated with specific receptor-mediated pathways (34). To determine whether these pathways are responsible for our results in epithelial cells, Calu-3 were exposed to a maximal 160 μg/mL OxPC dose in the presence of putative OxPC receptor blockers (Fig 9A). For each of the PAFR (WEB-2086; p=0.1179), EP2 (PF-04418948; p =0.5317), CD36 (SSO; p=0.7256) and TLR2/4 receptors (SsnB; p=0.1664), blocking agents did not inhibit general ROS accumulation. In contrast, the antioxidant NAC inhibited ROS in a dose-dependent manner, completely reversing ROS accumulation at a supraphysiological 10 mM dose (p=0.0004).

**Figure 9.**
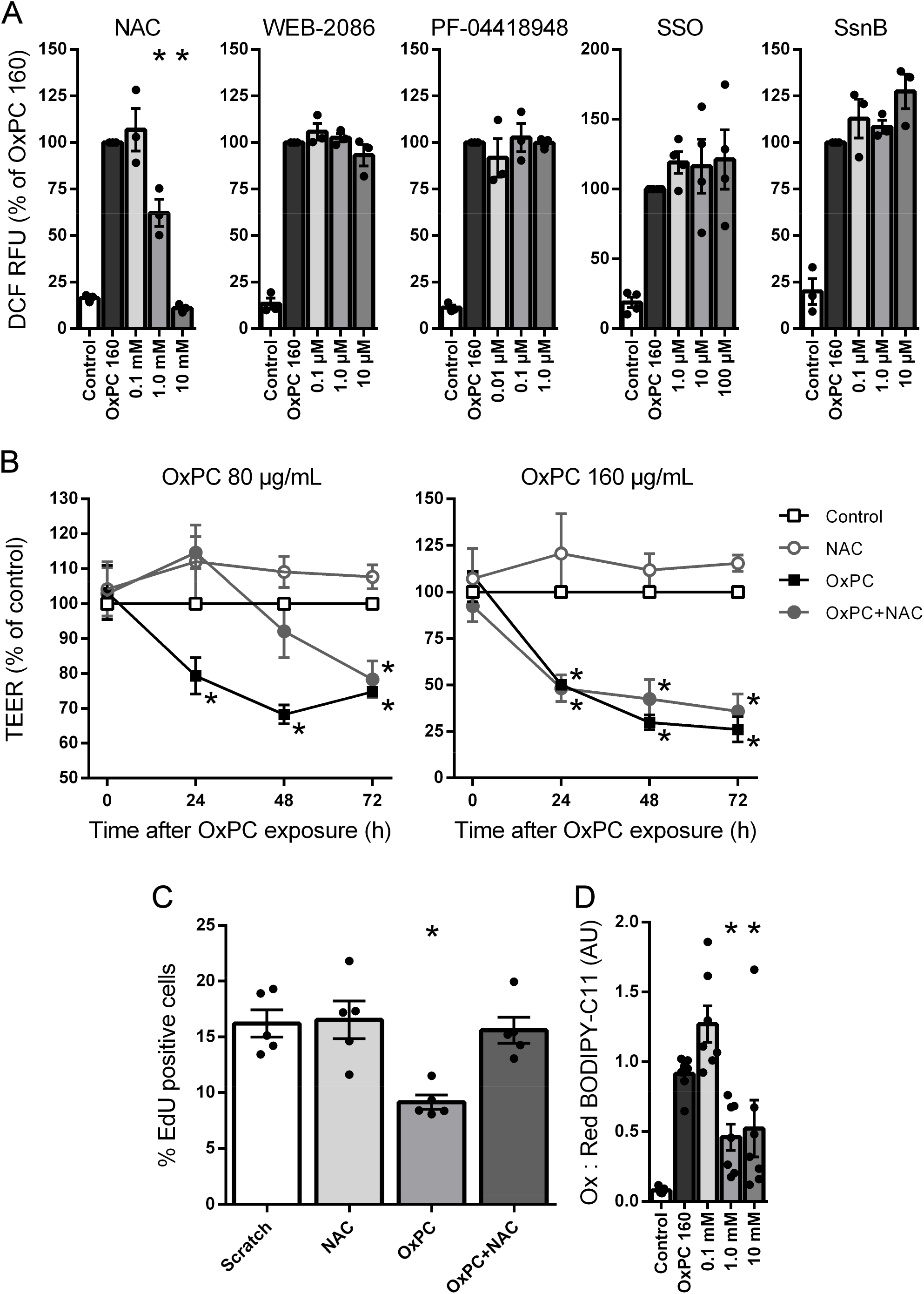
General ROS accumulation induced by OxPC in Calu-3 cells could be abrogated by the antioxidant NAC (A; p=0.0004, n=3). However, inhibitors of established OxPC receptors PAFR (WEB-2086; p=0.1179, n=3), EP2 (PF-04418948; p =0.5317, n=3), CD36 (SSO; p=0.7256, n=4) and TLR2/4 (SsnB; p=0.1664, n=3) each had no effect. Subsequently, 1 mM NAC was shown to temporarily delay barrier dysfunction induced by 80 μg/mL OxPC (B; interaction p=0.0003, time p=0.0013, OxPC p<0.0001, n=10) but not 160 μg/mL OxPC (interaction p<0.0001, time p<0.0001, OxPC p<0.0001, n=5). Similarly, the reduction in DNA synthesis observed with 80 μg/mL OxPC was not seen when co-treated with 1 mM NAC (C; p=0.0054, n=5), but lipid peroxidation persisted even at supramaximal NAC doses (D; p=0.0001, n=7). * represents significant difference (i.e., inhibition) from the OxPC 160 μg/mL condition (A and D) or from the 0 μg/mL OxPC control condition (B-D). Untreated Controls are shown for reference in A and D but are not included in the statistics.

### OxPC induced dysfunction can be partially prevented by NAC

To determine the capacity for NAC to prevent OxPC induced dysfunction, Calu-3 were treated concurrently with OxPC and NAC and re-assessed for barrier function, DNA synthesis after a wound, and lipid peroxidation. At a physiological 1 mM dose, NAC was able to delay the barrier dysfunction induced by 80 μg/mL OxPC for up to 48 hours (Fig 9B; OxPC p<0.0001). However, NAC could not abrogate the dysfunction induced by a maximal 160 μg/mL OxPC dose (OxPC p<0.0001). Consistent with these data, NAC prevented the reduction in DNA synthesis after a wound in the presence of 80 μg/mL OxPC (Fig 9D; p=0.0054), but could not completely inhibit lipid peroxidation to 160 μg/mL OxPC (Fig 9E; p=0.0001).

## DISCUSSION

### Epithelial dysfunction and physiological relevance

The airway epithelial barrier is the gateway to the lung and dysfunction of this barrier at a cellular level is associated with both asthma pathogenesis (29) and asthma severity (38). For this reason, we sought to explore the ability of OxPC to induce epithelial barrier dysfunction in the context of asthma. Our results indicate that OxPC can disrupt normal airway epithelial cell function and promote pathologic features similar to those seen in the airways of people with asthma. This includes a loss of barrier integrity (25), impaired capacity to re-establish barrier function after a wound (52), and further production of reactive oxygen species (14). This wide range of cellular dysfunction was accompanied by disruption of mitochondrial metabolism. Critically, each deleterious response was observed at doses below the threshold for detecting cell stress and loss of viability, i.e., 80 μg/mL, which is comparable to the level of OxPC seen in the airways of mouse models of asthma (31).

Wounding the epithelial layer increased the barrier-disrupting potency of OxPC, making OxPC effective at lower doses (Calu-3) and at earlier timepoints (ALI-nHBE). This suggests that OxPC could be important in potentiating the negative effects of allergenic insults or other stressors in diseased airways. In this regard, exposing epithelial cells to HDM resulted in an increase in ROS that was considerably smaller than the level induced by low doses of OxPC. Additionally, OxPC exposure could stimulate further production of lipid peroxides indicating the potential for a self-propagating cycle that starts with allergen exposure. Interestingly, the negative effects of OxPC could be reversed by simply washing the cells suggesting they do not cause permanent cell damage or irrecoverable activation of cell death pathways, and intracellular accumulation is not required. Instead, dysfunction seems to be dependent only on overall OxPC exposure. Therefore, OxPC are potentially important molecules in the pathogenesis of asthma and interventions aimed at limiting their production or promoting their removal from the airway milieu could be novel therapeutic options for this disease.

OxPC have been identified as a bioactive lipid molecule relevant to many chronic human diseases. In atherosclerosis, blocking the action of OxPC with the antibody E06 in mice reduced the amount of atherosclerosis and aortic valve calcifications (44). Following renal ischemia reperfusion injury in rats, there was a significant elevation in 55 OxPC species which correlated with the severity of the injury (50). Recently, members of our team have identified OxPC in the airways of mice and humans following allergen challenge and these were shown to be correlated with methacholine responsiveness (41). In airway smooth muscle cells, OxPC promote an inflammatory phenotype, characterized by secretion of IL-6, IL-8, GM-CSF, the production of eicosanoids, and were controlled by signaling events through protein kinase C. We were unable to detect similar OxPC-linked inflammatory mediator secretion from Calu-3 cells (data not shown), suggesting that OxPC contribute to disease pathology through more than one distinct mechanism. Thus, developing treatments that target the actions of OxPCs in the airways could offer us a better way to ameliorate the effects of oxidative stress in asthma, therefore augmenting current treatment regimes.

### Mechanisms of oxidative stress and lipid peroxidation

OxPC have been linked to signaling events involving PAFR, EP2, CD36 and TLR2/4 (34, 57). We were unable to show that any of these receptors were involved in mediating accumulation of ROS in response to OxPC in this study; only pre-incubation with the antioxidant NAC was able to reduce the cellular accumulation of ROS. Subsequently, we showed that NAC co-treatment reduced lipid peroxidation, increased proliferation capacity after a wound, and delayed barrier dysfunction to a submaximal 80 μg/mL OxPC dose. NAC operates as a cysteine source for the production of glutathione (47) and the glutathione anti-oxidant system is finely tuned for controlling cellular levels of lipid peroxides (60). In children with severe asthma, oxidized glutathione levels are significantly elevated and reduced glutathione levels are significantly repressed (18) indicating an elevated level of oxidative stress. Additionally, single nucleotide polymorphisms (SNPs) associated with increased asthma risk are present in some of the genes for glutathione-s-transferase enzymes, (32, 53). Disequilibrium in the glutathione system could thus indicate a diminished ability to detoxify OxPC in the airways and may render people with asthma particularly susceptible to the effects of OxPC. An alternative antioxidant mechanism for NAC involves triggering intracellular sulfane sulfur production which has antioxidant and cytoprotective functions (15). In this regard, treatments that increase endogenous production of sulfane sulfur could be useful in decreasing the antioxidant burden in asthma, specifically related to OxPC.

Yet despite the theoretical benefits of antioxidant treatments for asthma, results from animal and patient studies are mixed. With the exception of particular vitamin E isoforms that may specifically detoxify lipid peroxides (27, 43), dietary antioxidant supplementation seems to provide minimal benefit beyond a decrease in exhaled FeNO (1, 42, 54, 59). This may be consistent with our observations that NAC was unable to completely reverse lipid peroxidation and barrier dysfunction at a maximal 160 μg/mL OxPC dose, and that neither mitochondrial nor cytosolic superoxide are credible sources for the large magnitude of dysfunction we observed. This raises the possibility that the primary pathology induced by OxPC is specifically associated with lipid metabolism. Xenobiotic metabolism involving cytochrome P450, lipoxygenase and cyclooxygenase are well known to produce lipid peroxide by-products and these reactions can be self-propagating in the presence of oxygen (2). However, our demonstration of mitochondrial dysfunction after OxPC exposure, combined with previous observations that cardiolipin oxidation leads to lipid peroxidation in endothelial cells, suggests direct interference with mitochondrial lipid metabolism (6, 39). In this case, ROS and lipid peroxides would be produced that were detectable by our assays, but without necessarily producing a detectable superoxide intermediate. Metabolomic analysis of OxPC affected cells will be required to precisely determine which lipid peroxides are produced, the mitochondrial reactions/enzymes that are responsible, and how this process could be mitigated.

### Differential cell responses

Intriguingly, we found significant differences in the responsiveness of different epithelial cells to OxPC exposure. This may represent a culturing artifact or altered metabolic requirements of undifferentiated epithelial cells and cell lines but may also highlight how OxPC become disease relevant. Airway epithelium from people with asthma exhibits an elevated number of basal cells and a reduction in the number of cells expressing E-cadherin (24) suggesting a more immature phenotype. In this regard, airway epithelium from asthma may be more susceptible to the effects of OxPC due to its relatively undifferentiated state, although it is important to note that the cells used in this study are not sourced from asthmatic patients. Alternatively, commercial supplements for culturing primary epithelial cells in ALI contain hydrocortisone which has been shown to inhibit the inflammatory response of cultured epithelial cells (17) and support mucociliary transport (64). Importantly, glucocorticoids have been shown to increase the activity of antioxidant enzymes in asthma (11). We did not remove hydrocortisone from the media prior to experimentation which may have conferred some protection and reduced the effect sizes seen in primary cells. Finally, OxPC had a significant impact on DNA synthesis in our experiments suggesting that they may directly inhibit proliferation. Primary cells are relatively quiescent compared with the cell lines used in this study perhaps indicating that proliferating cells, or those that have an associated high metabolic requirement, are particularly susceptible to OxPC exposure. It will therefore be important to study how prolonged OxPC treatment alters the growth and differentiation trajectory of primary epithelial cells, to determine the effect on mucociliary phenotype and barrier function.

### Summary

We have demonstrated that OxPC induce dose-dependent cellular dysfunction that may potentiate other stressors to manifest as impaired epithelial barrier function in diseased airways. Future work will need to ascertain whether the cellular dysfunction we have observed is a product of the combination and total quantity of OxPC applied or can be attributable to a single OxPC species operating through a specific mechanism. Understanding the role of individual OxPC in causing epithelial cell dysfunction will help us to identify potential treatment options that protect the cell or ‘mop-up’ OxPC in a more specific manner than NAC, and more completely prevent lipid peroxide accumulation and barrier dysfunction. It will also be important to ascertain whether cells from asthmatic patients might be more susceptible to OxPC because of deficiencies in antioxidant systems or immaturity of epithelial phenotype. Finally, by using mice that express the anti-OxPC antibody (E06), and determining whether OxPC can augment the pathology associated with HDM exposure, it will be possible to discover whether OxPC are sufficient and necessary for the pathogenesis of airway disease.

## ACKNOWLEDGEMENTS

Adrian West is supported by NSERC Discovery Grant # RGPIN-2014-06412 and Research Manitoba New Investigator Operating Grant # 1653. Chris Pascoe is supported by Research Manitoba Postdoctoral Fellowship # 1772, a Canadian Respiratory Research Network Fellowship and the Canadian Institutes of Health Research Banting Postdoctoral Fellowship. Neilloy Roy is supported by Research Manitoba and CHRIM Graduate Studentship # 1329. Jignesh Vaghasiya is supported by Research Manitoba and CHRIM Graduate Studentship # #3735 and #4196. Andrew Halayko is supported by the Canada Research Chairs Program, and portions of this work were supported by funding from the CIHR Canadian Respiratory Research Network and the Children’s Hospital Research Institute of Manitoba. The authors would like to thank Gerald Stelmack for assistance with Beas-2B cell culture.

## AUTHOR CONTRIBUTIONS

Chris Pascoe co-wrote the manuscript, designed experiments, generated OxPC, and performed cell culture, ROS/superoxide assays, lipid peroxidation assays, and mitochondrial function assays.

Neilloy Roy performed paracellular permeability assays.

Emily Turner-Brannen performed cell culture, cell stress/viability assays, TEER measurements, mRNA abundance analysis, proliferation assays, and mitochondrial function assays.

Alexander Schultz performed HDM ROS and HDM lipid peroxide assays.

Jignesh Vaghasiya generated OxPC.

Amir Ravandi conceived methods to generate OxPC, and performed mass spectrometric assessment of the OxPC mixture.

Andrew Halayko helped conceive the study and in vitro protocols for using OxPC, edited the manuscript, generated OxPC, and supplied OxPC receptor inhibitors.

Adrian West conceived the study, co-wrote the manuscript, designed/supervized all aspects of experimentation, and performed data/statistical analysis, cell culture and TEER measurements.

